# Comparative Genomic Analysis of Rapidly Evolving SARS-CoV-2 Viruses Reveal Mosaic Pattern of Phylogeographical Distribution

**DOI:** 10.1101/2020.03.25.006213

**Authors:** Roshan Kumar, Helianthous Verma, Nirjara Singhvi, Utkarsh Sood, Vipin Gupta, Mona Singh, Rashmi Kumari, Princy Hira, Shekhar Nagar, Chandni Talwar, Namita Nayyar, Shailly Anand, Charu Dogra Rawat, Mansi Verma, Ram Krishan Negi, Yogendra Singh, Rup Lal

## Abstract

The Coronavirus Disease-2019 (COVID-19) that started in Wuhan, China in December 2019 has spread worldwide emerging as a global pandemic. The severe respiratory pneumonia caused by the novel SARS-CoV-2 has so far claimed more than 60,000 lives and has impacted human lives worldwide. However, as the novel SARS-CoV-2 displays high transmission rates, their underlying genomic severity is required to be fully understood. We studied the complete genomes of 95 SARS-CoV-2 strains from different geographical regions worldwide to uncover the pattern of the spread of the virus. We show that there is no direct transmission pattern of the virus among neighboring countries suggesting that the outbreak is a result of travel of infected humans to different countries. We revealed unique single nucleotide polymorphisms (SNPs) in nsp13-16 (ORF1b polyprotein) and S-Protein within 10 viral isolates from the USA. These viral proteins are involved in RNA replication, indicating highly evolved viral strains circulating in the population of USA than other countries. Furthermore, we found an amino acid addition in nsp16 (mRNA cap-1 methyltransferase) of the USA isolate (MT188341) leading to shift in amino acid frame from position 2540 onwards. Through the construction of SARS-CoV-2-human interactome, we further revealed that multiple host proteins (PHB, PPP1CA, TGF-β, SOCS3, STAT3, JAK1/2, SMAD3, BCL2, CAV1 & SPECC1) are manipulated by the viral proteins (nsp2, PL-PRO, N-protein, ORF7a, M-S-ORF3a complex, nsp7-nsp8-nsp9-RdRp complex) for mediating host immune evasion. Thus, the replicative machinery of SARS-CoV-2 is fast evolving to evade host challenges which need to be considered for developing effective treatment strategies.

## Background

Since the current outbreak of pandemic coronavirus disease 2019 (COVID-19) caused by Severe Acute Respiratory Syndrome-related Coronavirus-2 (SARS-CoV-2), the assessment of the biogeographical pattern of SARS-CoV-2 isolates and the mutations at nucleotide and protein level is of high interest to many research groups [1, 2, 3]. Coronaviruses (CoVs), members of *Coronaviridae* family, order *Nidovirales*, have been known as human pathogens from the last six decades [4]. Their target is not just limited to humans, but also other mammals and birds [5]. Coronaviruses have been classified under alpha, beta, gamma and delta-coronavirus groups [6] in which former two are known to infect mammals while the latter two primarily infect bird species [7]. Symptoms in humans vary from common cold to respiratory and gastrointestinal distress of varying intensities. In the past, more severe forms caused major outbreaks that include Severe Acute Respiratory Syndrome (SARS-CoV) (outbreak in 2003, China) and Middle East Respiratory Syndrome (MERS-CoV) (outbreak in 2012, Middle East) [8]. Bats are known to host coronaviruses acting as their natural reservoirs which may be transmitted to humans through an intermediate host. SARS-CoV and MERS-CoV were transmitted from intermediate hosts, palm civets and camel, respectively [9, 10]. It is not, however, yet clear which animal served as the intermediate host for transmission of SARS-CoV-2 transmission from bats to humans which is most likely suggested to be a warm-blooded vertebrate [11].

The inherently high recombination frequency and mutation rates of coronavirus genomes allow for their easy transmission among different hosts. Structurally, they are positive-sense single stranded RNA (ssRNA) virions with characteristic spikes projecting from the surface of capsid coating [12, 13]. The spherical capsid and spikes give them crown-like appearance due to which they were named as ‘corona’, meaning ‘crown’ or ‘halo’ in *Latin*. Their genome is nearly 30 Kb long, largest among the RNA viruses, with 5’cap and 3’ polyA tail, for translation [14]. Coronavirus consists of four main proteins, spike (S), membrane (M), envelope (E) and nucleocapsid (N). The spike (∼150 kDa) mediates its attachment to host receptor proteins [15]. Membrane protein (∼25-30 kDa) attaches with nucleocapsid and maintains curvature of virus membrane [16]. E protein (8-12 kDa) is responsible for the pathogenesis of the virus as it eases assembly and release of virion particles and also has ion channel activity as integral membrane protein [17]. N-protein, the fourth protein, helps in the packaging of virus particles into capsids and promotes replicase-transcriptase complex (RTC) [18].

Recently, in December 2019, the outbreak of novel beta-coronavirus (2019-nCoV) or SARS-CoV-2 in Wuhan, China has shown devastating effects worldwide (https://www.who.int/docs/default-source/coronaviruse/situation-reports/20200403-sitrep-74-covid-19-mp.pdf?sfvrsn=4e043d03_4)).World Health Organization (WHO) has declared COVID-19, the disease caused by the novel SARS-CoV-2 a pandemic, affecting more than 186 countries and territories where USA has most reported cases 2,13,600 and Italy has highest mortality rate 12.08% (1,15,242 infected individuals, 13,917 deaths) (WHO situation report-74). As on date (April 4, 2020), more than 1 million individuals have been infected by SARS-CoV-2 and nearly 60,000 have died worldwide. Virtually, all human lives have been impacted with no foreseeable end of the pandemic. A recent study on ten novel coronavirus strains by Lu *et al.*, suggested that SARS-CoV-2 has sufficiently diverged from SARS-CoV [19]. SARS-CoV-2 is assumed to have originated from bats, which serve as a reservoir host of the virus [19]. A recent study has shown similar mutation patterns in Bat-SARS-CoV RaTG13 and SARS CoV-2, but the dataset was limited to 21 strains including few SARS-CoV-2 strains and other neighbors [20]. Other studies have also reported the genome composition and divergence patterns of SARS-CoV-2 [3, 21]. However, no study has yet explained the biogeographical pattern of this emerging pathogen. In this study, we selected 95 strains of SARS-CoV-2, isolated and sequenced from 11 different countries to understand the transmission patterns, evolution and pathogenesis of the virus. Using core genome and Single Nucleotide Polymorphism (SNP) based phylogeny, we attempted to uncover any existence of a transmission pattern of the virus across the affected countries, which was not known earlier. We analyzed the ORFs of the isolates to reveal unique point mutations and amino-acid substitutions/additions in the isolates from the USA. In addition, we analyzed the gene/protein mutations in these novel strains and estimated the direction of selection to decipher their evolutionary divergence rate. Further, we also established the interactome of SARS-CoV-2 with the human host proteins to predict the functional implications of the viral infection host cells. The results obtained from the analyses indicate the high severity of SARS-CoV-2 isolates with the inherent capability of unique mutations and the evolving viral replication strategies to adapt to human hosts.

## Materials and Methods

### Selection of genomes and annotation

Sequences of different strains were downloaded from NCBI database https://www.ncbi.nlm.nih.gov/genbank/sars-cov-2-seqs/ (Table 1). A total of 97 genomes were downloaded on March 19, 2020 from NCBI database and based on quality assessment two genomes with multiple Ns were removed from the study. Further the genomes were annotated using Prokka [22]. A manually annotated reference database was generated using the Genbank file of Severe acute respiratory syndrome coronavirus 2 isolate-SARS-CoV-2/SH01/human/2020/CHN (Accession number: MT121215) and open reading frames (ORFs) were predicted against the formatted database using prokka (-gcode 1) [22]. Further the GC content information was generated using QUAST standalone tool [23].

**Table 1:**
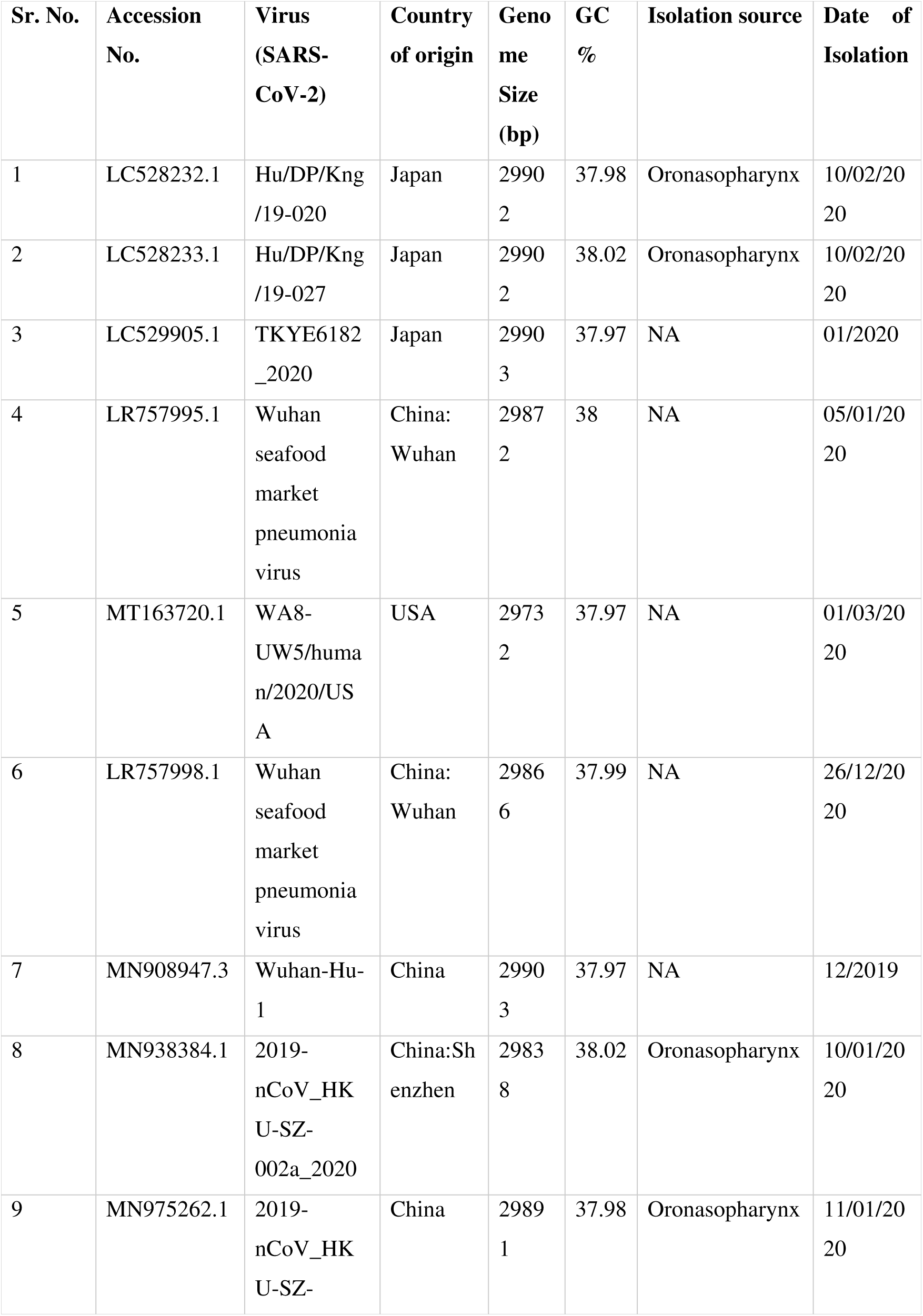
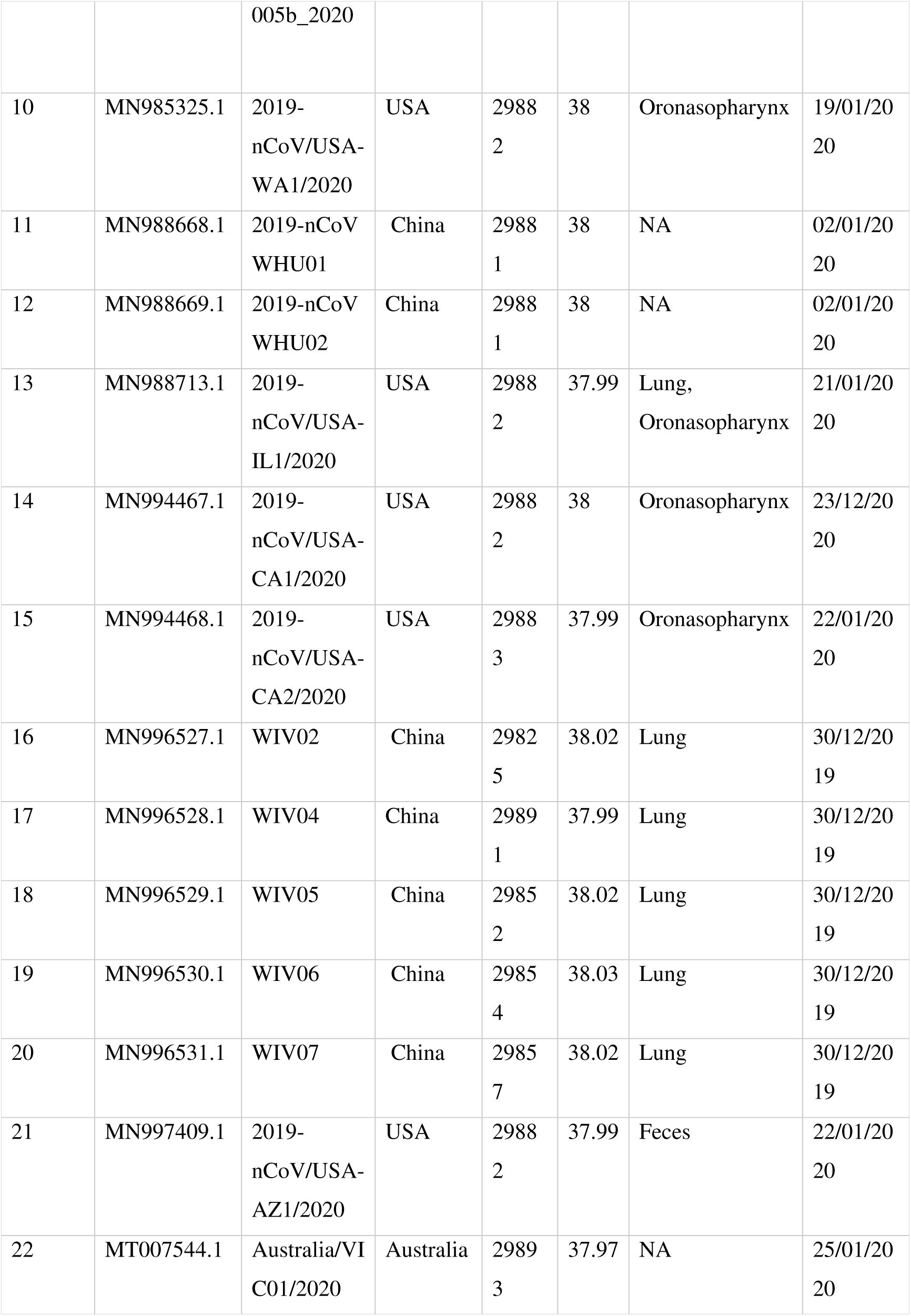
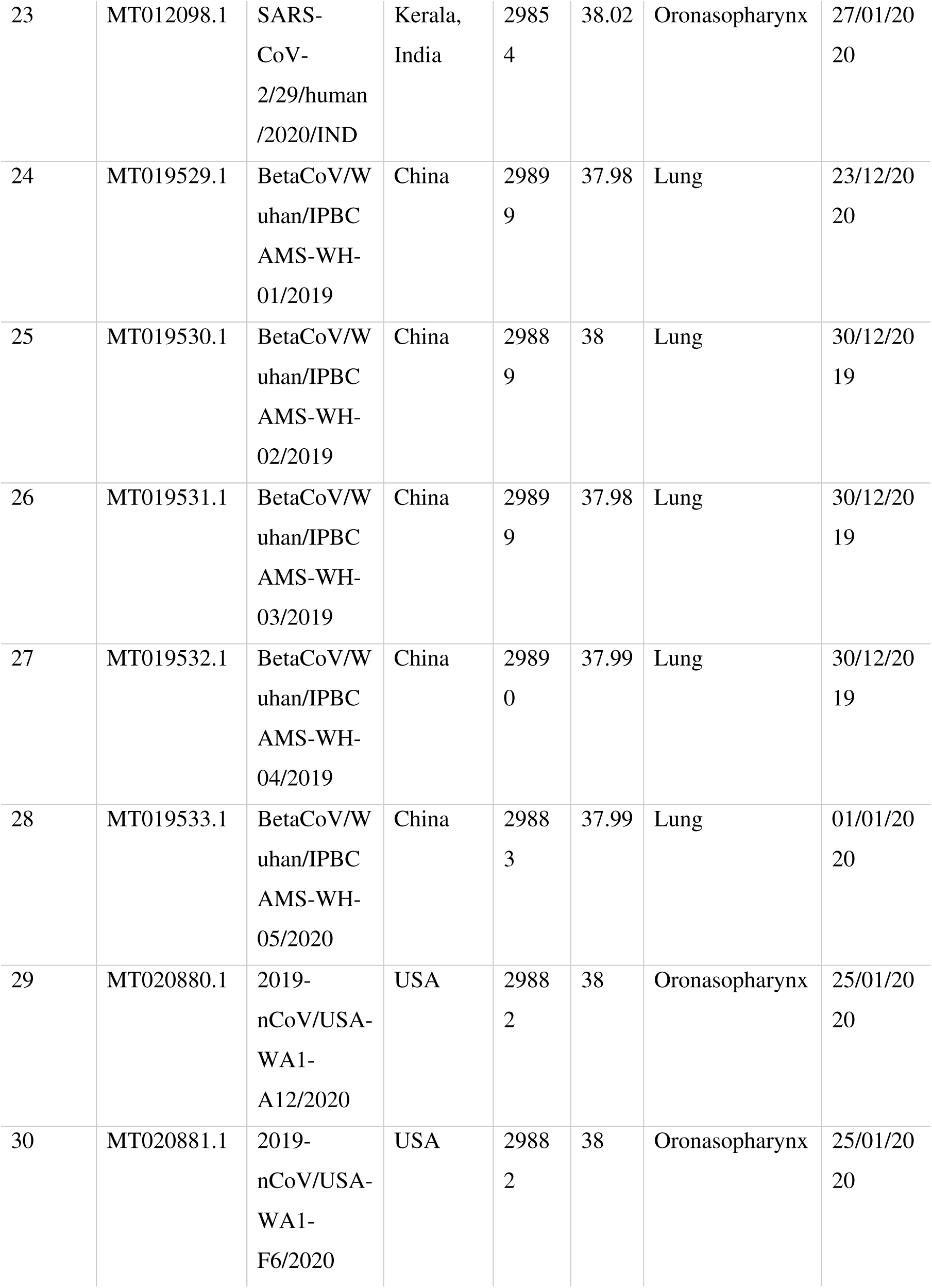
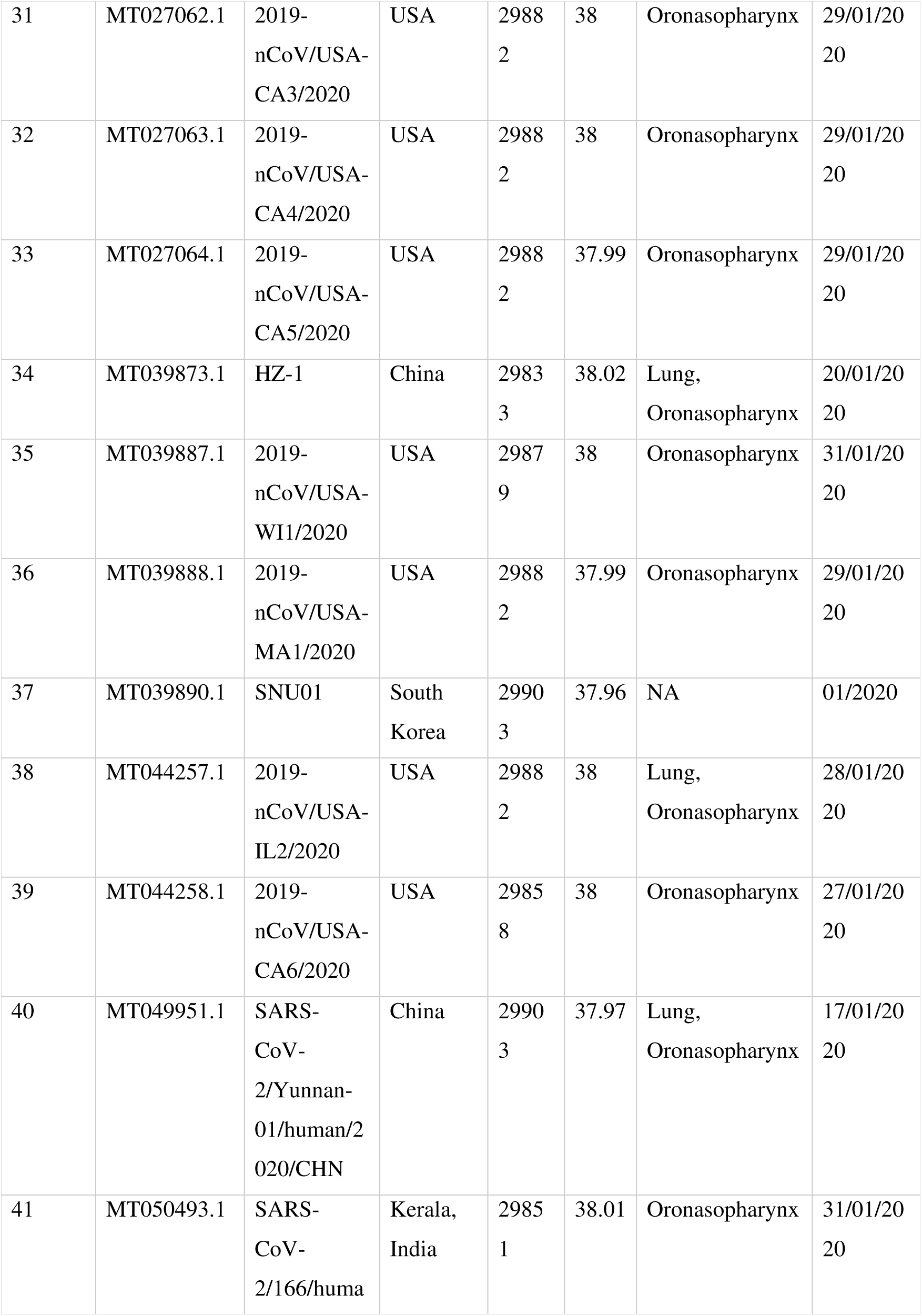
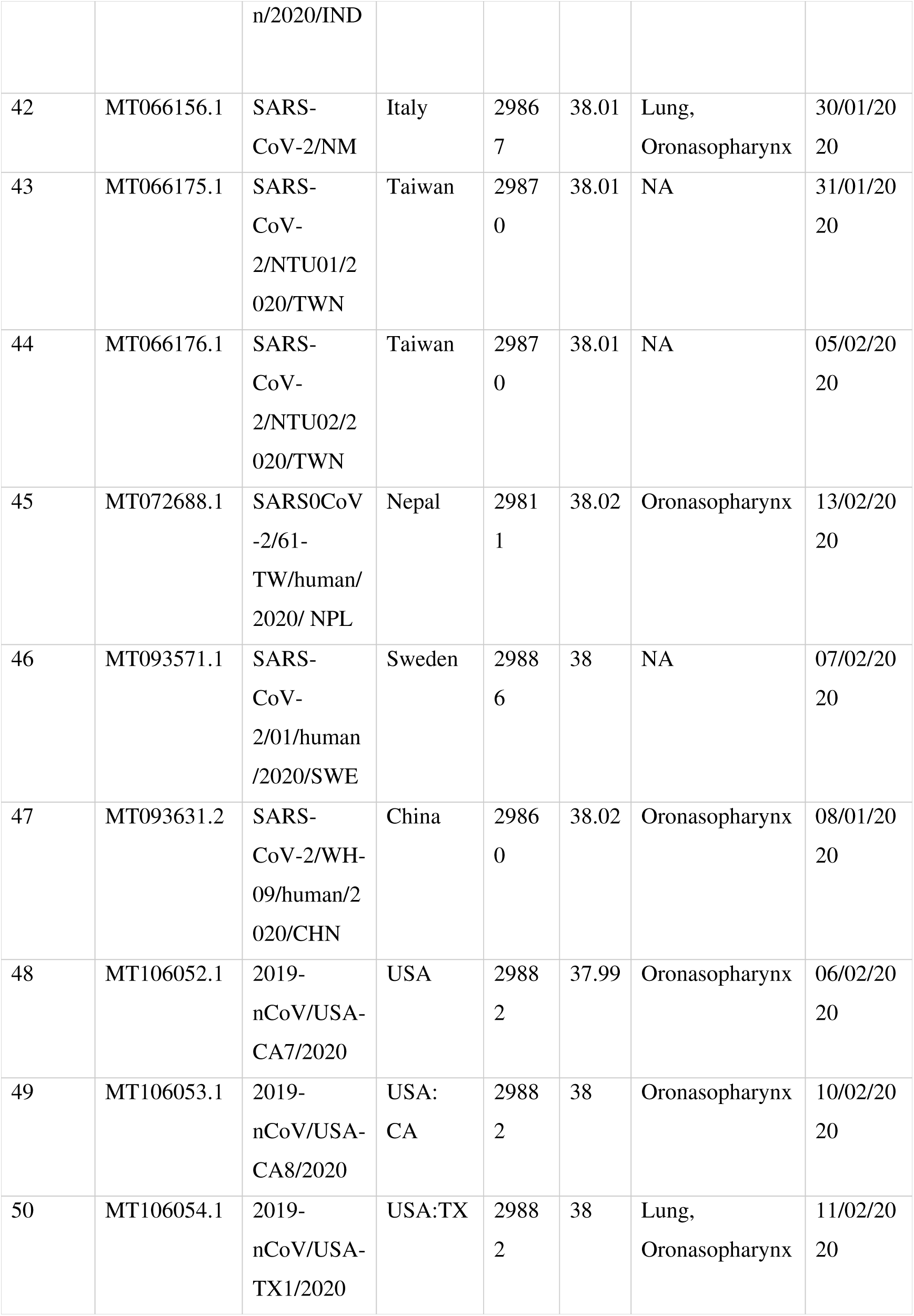
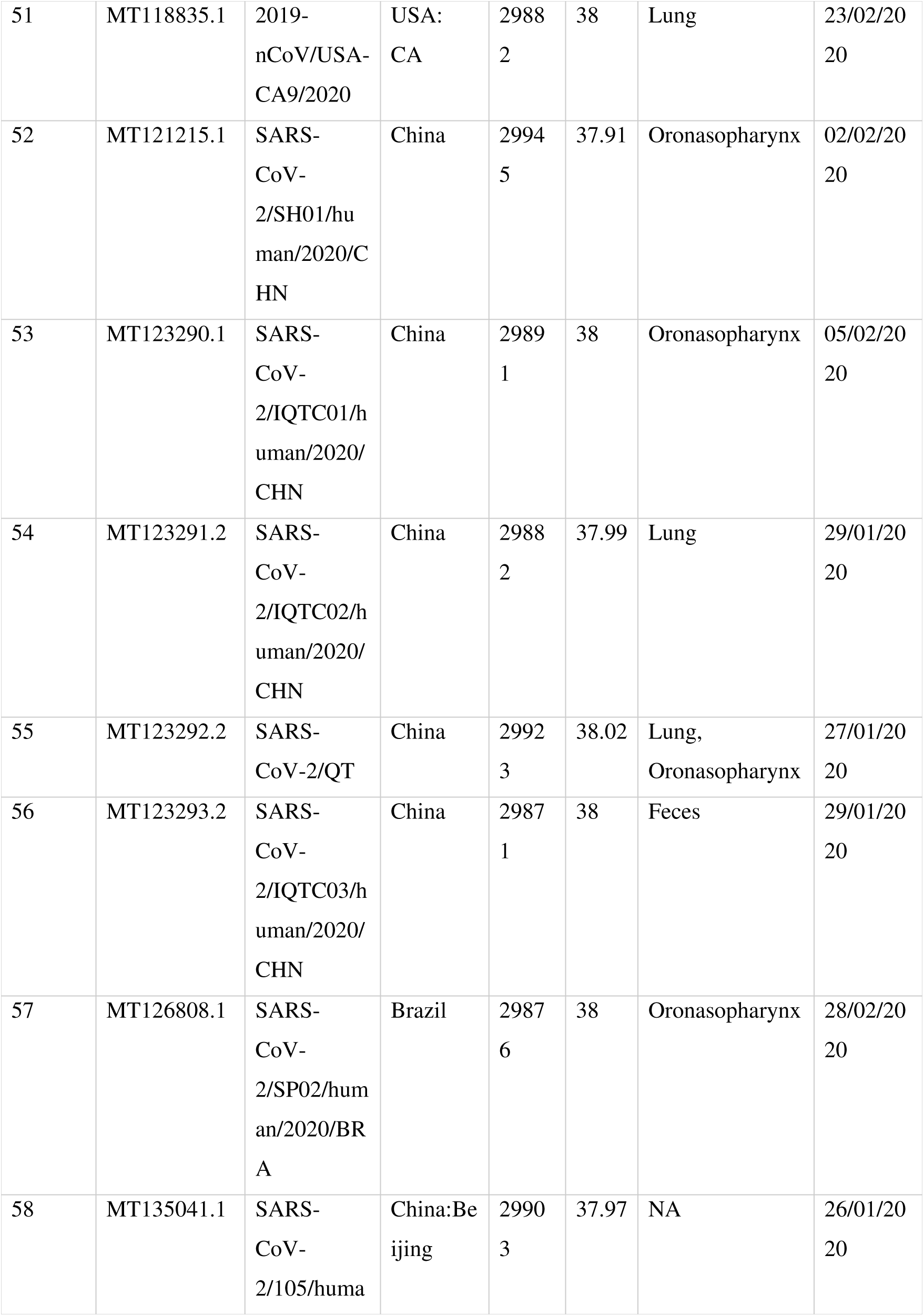
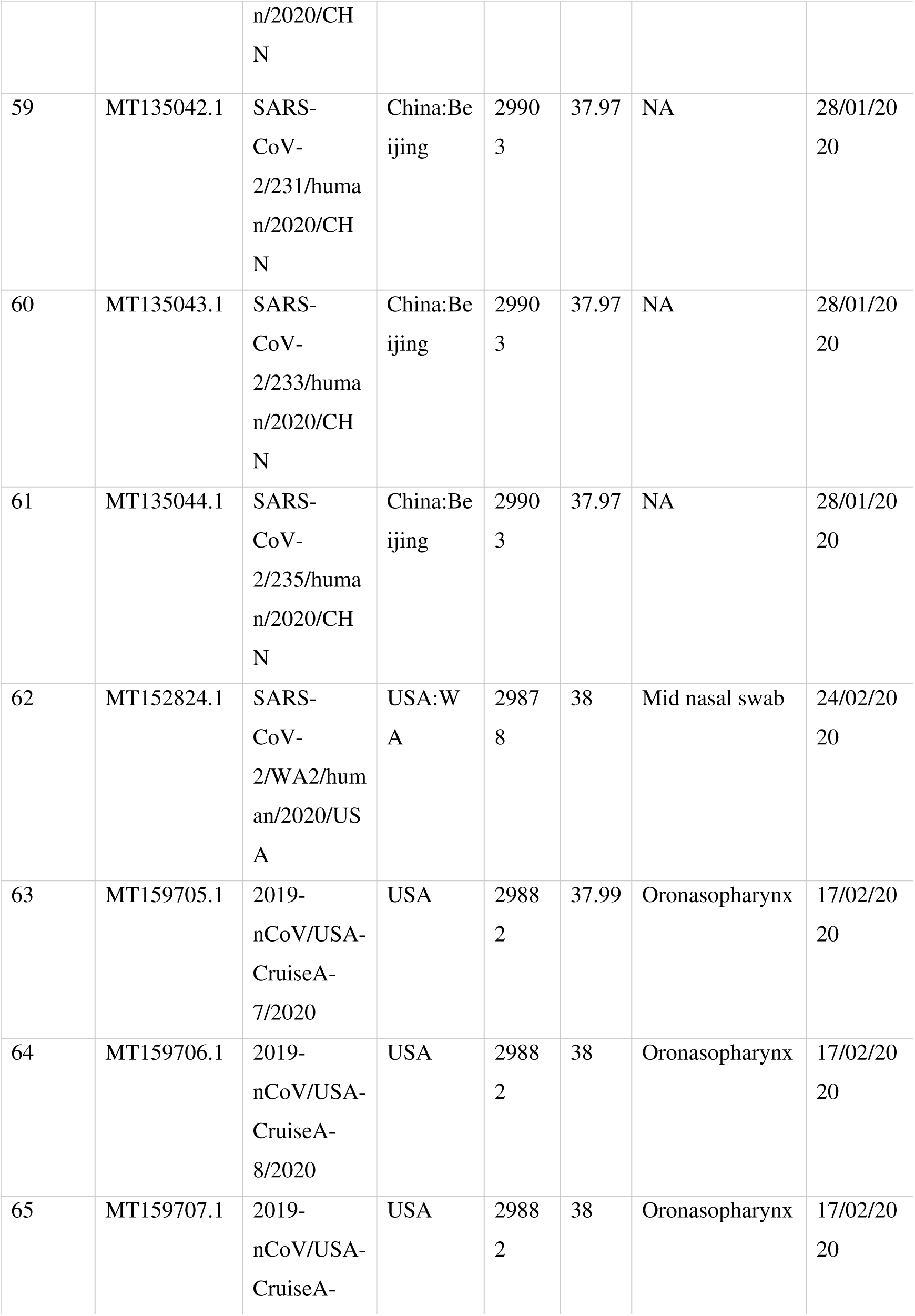
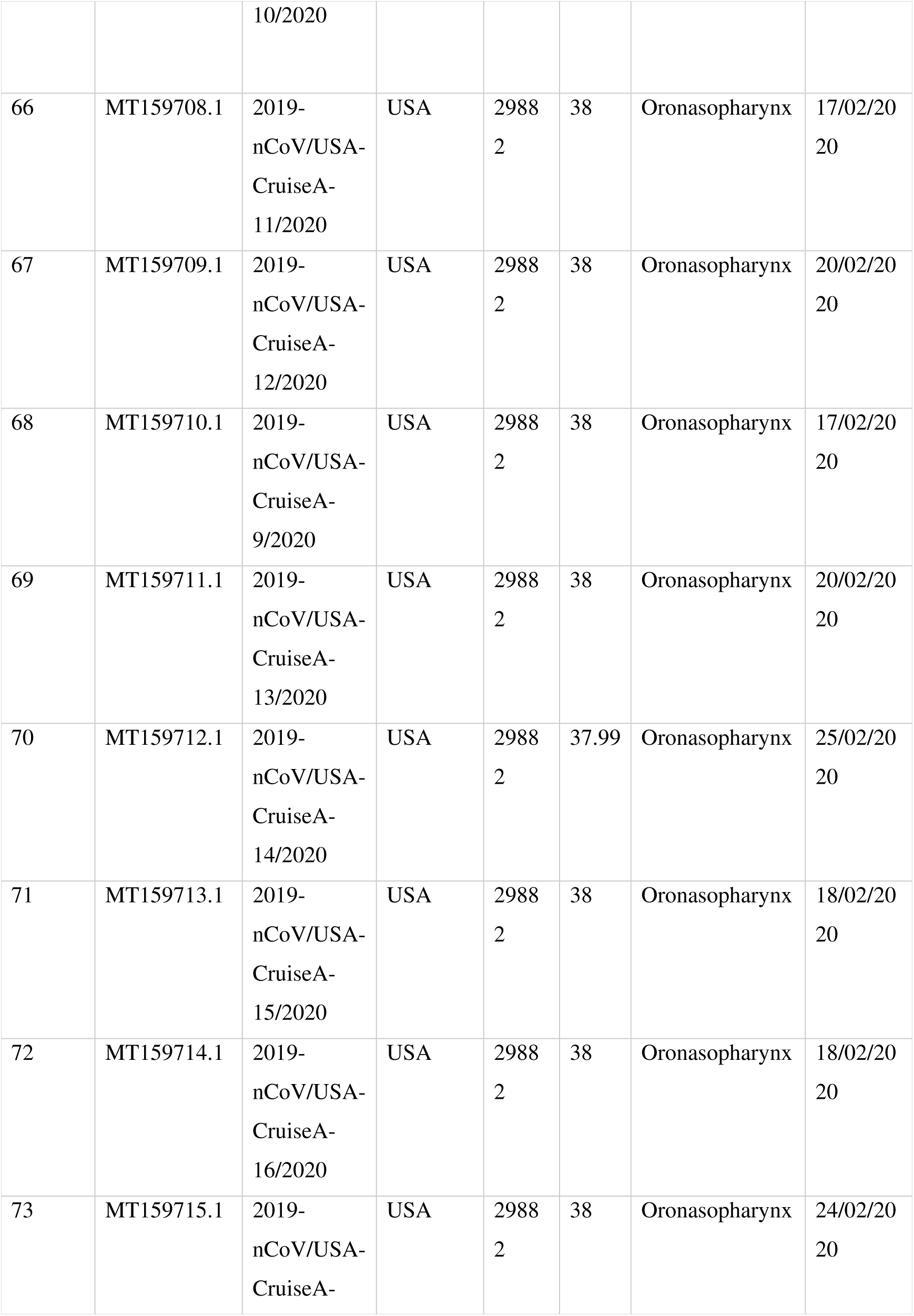
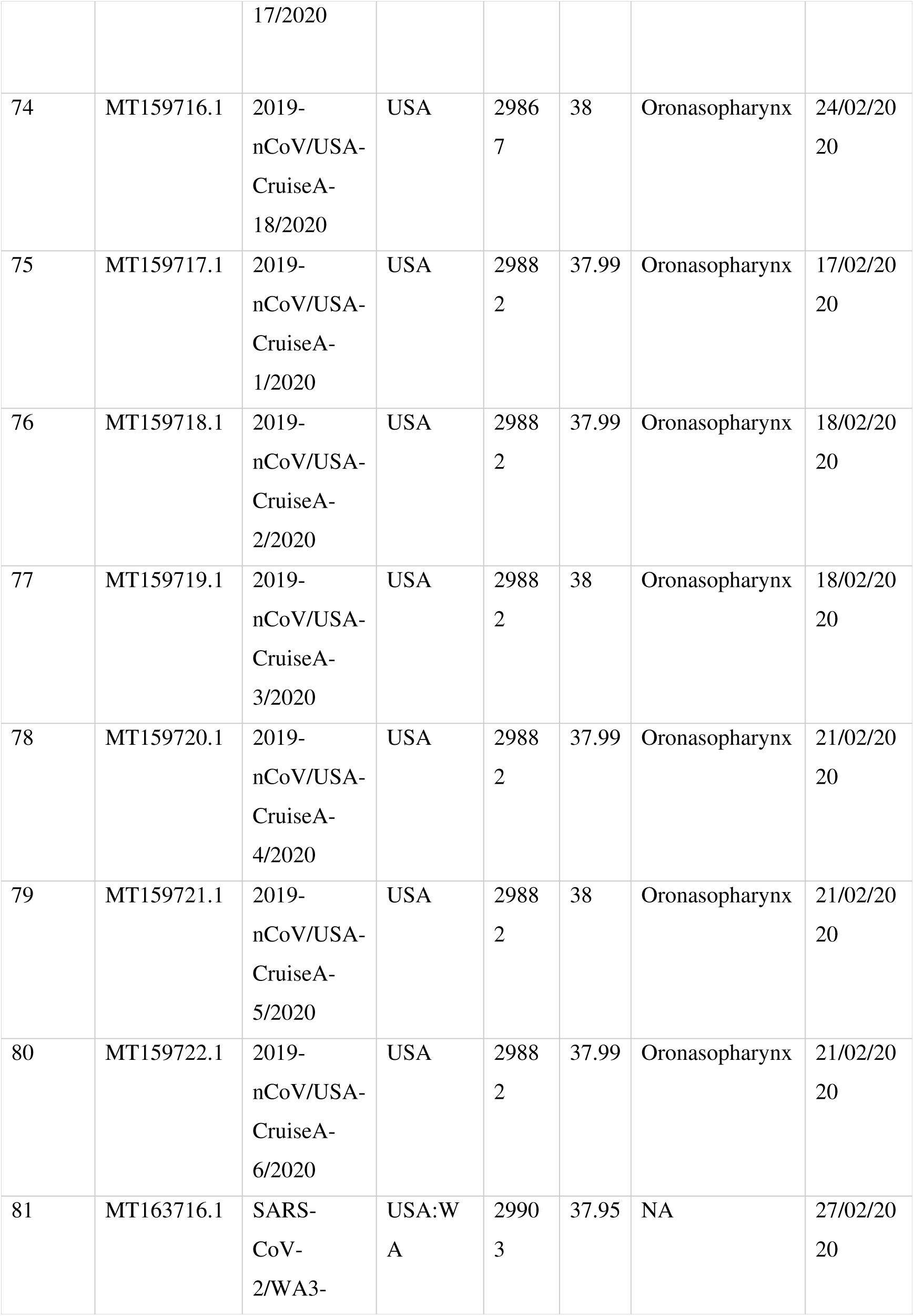
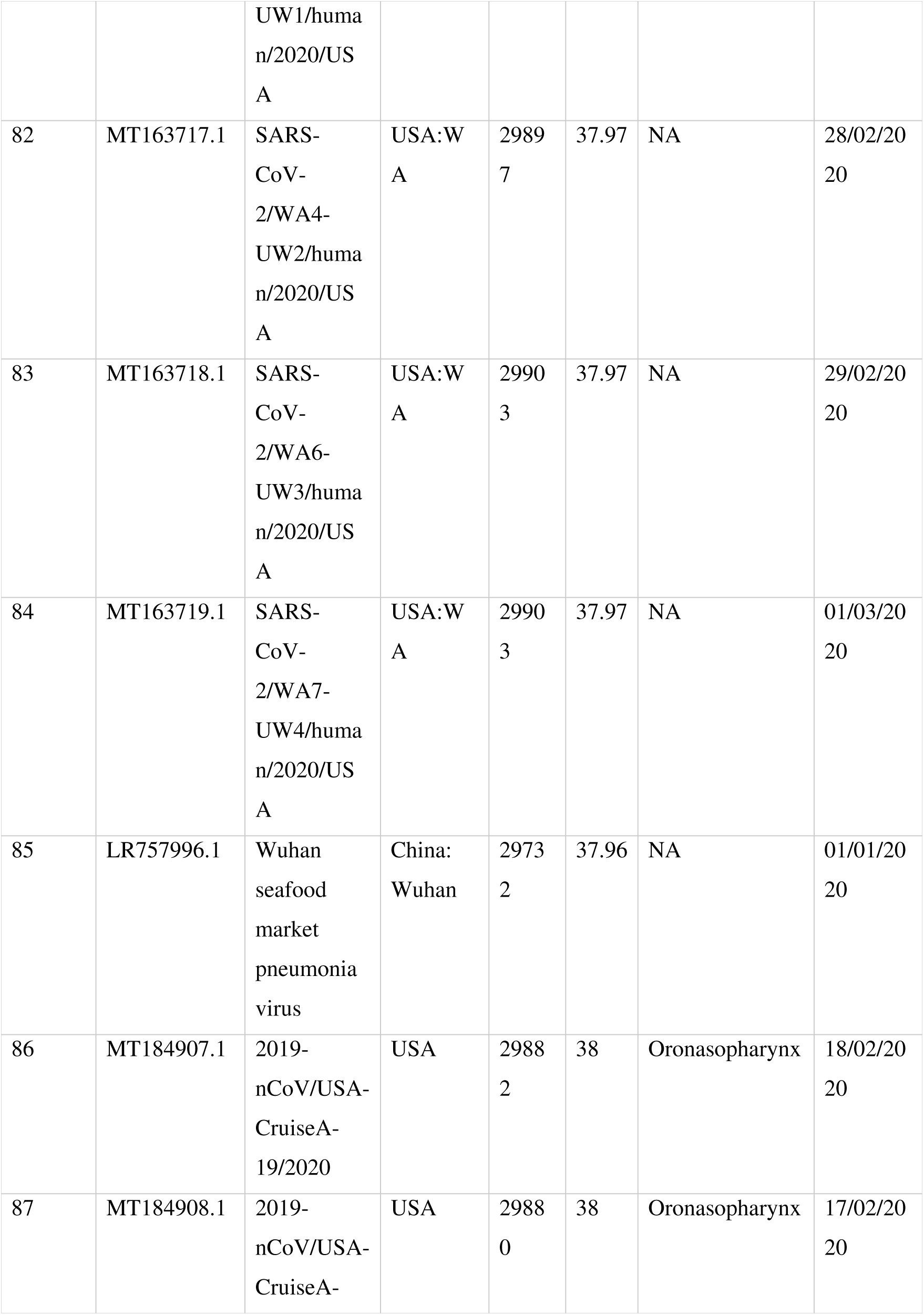
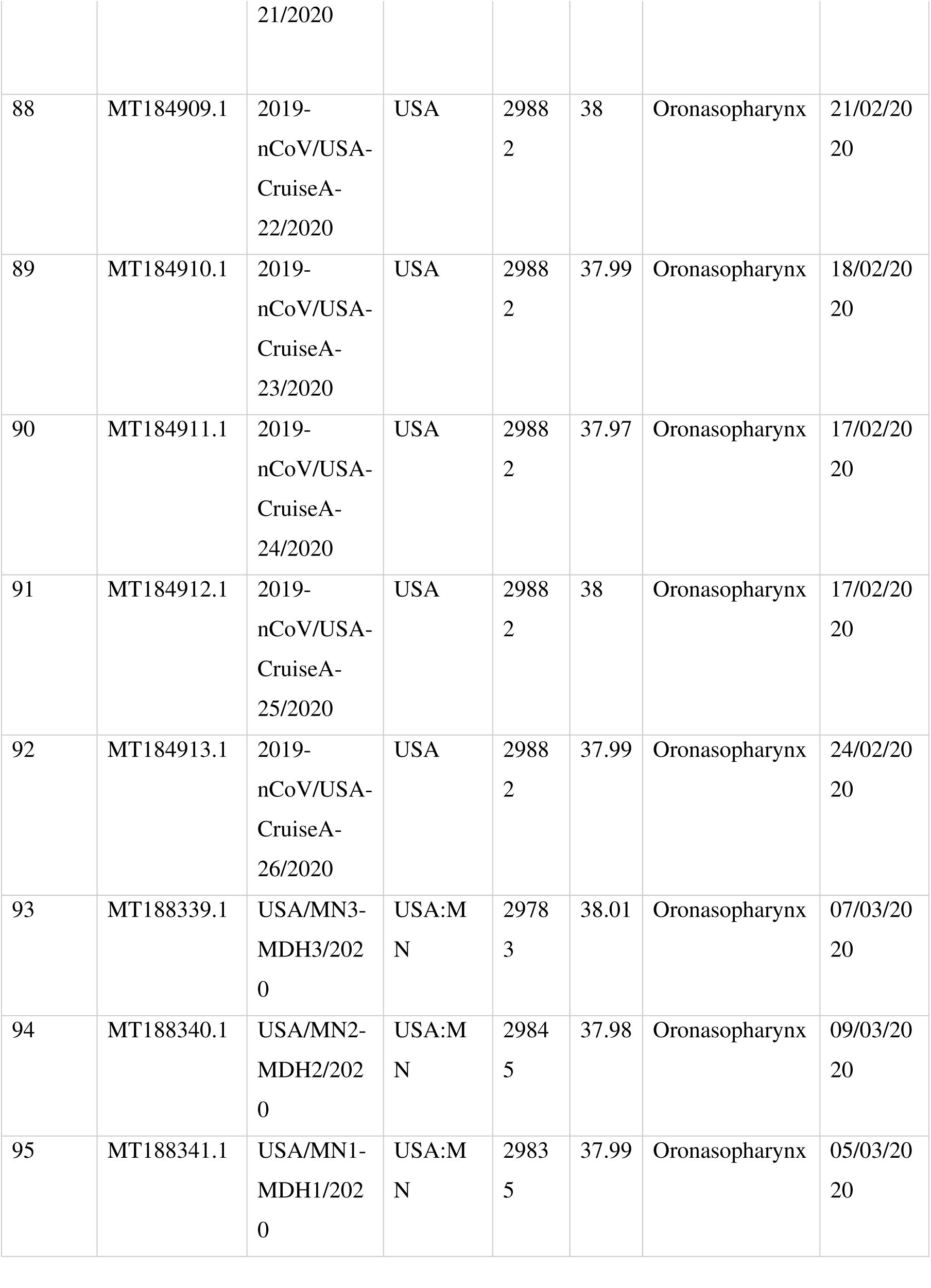
General genomic attributes of SARS-CoV-2 strains.

### Analysis of natural selection

To determine the evolutionary pressure on viral proteins, dN/dS values were calculated for 9 ORFs of all strains. The orthologous gene clusters were aligned using MUSCLE v3.8 [24] and further processed for removing stop codons using HyPhy v2.2.4 [25]. Single-Likelihood Ancestor Counting (SLAC) method in Datamonkey v2.0 [26] (http://www.datamonkey.org/slac) was used to calculate dN/dS value for each orthologous gene cluster. The dN/dS values were plotted in R (R Development Core Team, 2015).

### Phylogenetic analysis

To infer the phylogeny, the core gene alignment was generated using MAFFT [27] present within the Roary Package [28]. Further, the phylogeny was inferred using the Maximum Likelihood method based and Tamura-Nei model [29] in MEGAX [30] and visualized in interactive Tree of Life (iTOL) [31] and GrapeTree [32].

To determine the single nucleotide polymorphism (SNP), whole-genome alignments were made using libMUSCLE aligner. For this, we used annotated genbank of SARS-CoV-2/SH01/human/2020/CHN (Accession no. MT121215) as the reference in the parsnp tool of Harvest suite [33]. As only genomes within a specified MUMI distance threshold are recruited, we used option -c to force include all the strains. For output, it produced a core-genome alignment, variant calls and a phylogeny based on Single nucleotide polymorphisms. The SNPs were further visualized in Gingr, a dynamic visual platform [33]. Further, the tree was visualized in interactive Tree of Life (iTOL) [31].

### SARS-CoV-2 protein annotation and host-pathogenic interactions

SARS-CoV-2/SH01/human/2020/CHN virus genome having accession no. MT121215.1 was used for protein-protein network analysis. Since, none of the SARS-CoV-2 genomes are updated in any protein database, we first annotated the genes using BLASTp tool [34]. The similarity searches were performed against SARS-CoV isolate Tor2 having accession no. AY274119 selected from NCBI at default parameters. The annotated SARS-CoV-2 proteins were mapped against viruSITE [35] and interaction databases such as Virus.STRING v10.5 [36] and IntAct [37] for predicting their interaction against host proteins. These proteins were either the direct targets of HCoV proteins or were involved in critical pathways of HCoV infection identified by multiple experimental sources. To build a comprehensive list of human PPIs, we assembled data from a total of 18 bioinformatics and systems biology databases with five types of experimental evidence: (i) binary PPIs tested by high-throughput yeast two-hybrid (Y2H) systems; (ii) binary, physical PPIs from protein 3D structures; (iii) kinase-substrate interactions by literature-derived low-throughput or high-throughput experiments; (iv) signaling network by literature-derived low-throughput experiments; and (v) literature-curated PPIs identified by affinity purification followed by mass spectrometry (AP-MS), Y2H, or by literature-derived low [36, 38].

Filtered proteins (confidence value: 0.7) were mapped to their Entrez ID [39] based on the NCBI database used for interactome analysis. HPI were stimulated using Cytoscape v.3.7.2 [40].

### Functional enrichment analysis

Next, functional studies were performed using the Kyoto Encyclopedia of Genes and Genomes (KEGG) [41, 42] and Gene Ontology (GO) enrichment analyses using UniProt database [43] to evaluate the biological relevance and functional pathways of the HCoV-associated proteins. All functional analyses were performed using STRING enrichment and STRINGify, plugin of Cytoscape v.3.7.2 [40]. Network analysis was performed by tool NetworkAnalyzer, plugin of Cytoscape with the orthogonal layout.

## Results and Discussion

### General genomic attributes of SARS-CoV-2

In this study, we analyzed a total of 95 SARS-CoV-2 strains (available on March 19, 2020) isolated between December 2019-March 2020 from 11 different countries namely USA (n=52), China (n=30), Japan (n=3), India (n=2), Taiwan (n=2) and one each from Australia, Brazil, Italy, Nepal, South Korea and Sweden. A total of 68 strains were isolated from either oronasopharynges or lungs, while two of them were isolated from faeces suggesting both respiratory and gastrointestinal connection of SARS-CoV-2 (Table 1). No information of the source of isolation of the remaining isolates is available. The average genome size and GC content were found to be 29879 ± 26.6 bp and 37.99 ± 0.018%, respectively. All these isolates were found to harbor 9 open reading frames coding for ORF1a (13218 bp) and ORF1b (7788 bp) polyproteins, surface glycoprotein or S-protein (3822 bp), ORF3a protein (828 bp), membrane glycoprotein or M-protein (669 bp), ORF6 protein (186 bp), ORF7a protein (366 bp), ORF8 protein (366 bp), and nucleocapsid phosphoprotein or N-protein (1260 bp) which agrees with a recently published study [44]. The ORF1a harbors 12 non-structural protein (nsp) namely nsp1, nsp2, nsp3 (papain-like protease or PLpro domain), nsp4, nsp5 (3C-like protease or 3CLpro), nsp6, nsp7, nsp8, nsp9, nsp10, nsp11and nsp12 (RNA-dependent RNA polymerase or RdRp) [44]. Similarly, ORF1b contains four putative nsp’s namely nsp13 (helicase or Hel), nsp14 (3′-to-5′ exoribonuclease or ExoN), nsp15 and nsp16 (mRNA cap-1 methyltransferase).

### Phylogenomic analysis: defining evolutionary relatedness

Our analysis revealed that strains of human infecting SARS-CoV-2 are novel and highly identical (>99.9%). A recent study established the closest neighbor of SARS-CoV-2 as SARSr-CoV-RaTG13, a bat coronavirus [45]. As COVID19 transits from epidemic to pandemic due to extremely contagious nature of the SARS-CoV-2, it was interesting to draw the relation between strains and their geographical locations. In this study, we employed two methods to delineate phylogenomic relatedness of the isolates: core genome (Figure 1A & C) and single nucleotide polymorphisms (SNPs) (Figure 1B). Phylogenies obtained were annotated with country of isolation of each strain (Figure 1A & B). The phylogenetic clustering was found majorly concordant by both core-genome (Figure 1A) and SNP based methods (Figure 1B). The strains formed a monophyletic clade, in which MT093571.1 (South Korea) and MT039890.1 (Sweden) were most diverged. Focusing on the edge-connection between the neighboring countries from where the transmission is more likely to occur, we noted a strain from Taiwan (MT066176) closely clustered with another from China (MT121215.1). With the exception of these two strains, we did not find any connection between strains of neighboring countries. Thus, most strains belonging to the same country clustered distantly from each other and showed relatedness with strains isolated from distant geographical locations (Figure 1A & B). For instance, a SARS-CoV-2 strain isolated from Nepal (MT072688) clustered with a strain from USA (MT039888). Also, strains from Wuhan (LR757998 and LR757995), where the virus was originated, showed highest identity with USA as well as China strains; strains from India, MT012098 and MT050493 clustered closely with China and USA strains, respectively (Figure 1A & B). Similarly, Australian strain (MT007544) showed close clustering with USA strain (Figure 1A & B) and one strain from Taiwan (MT066175) clustered nearly with Chinese isolates (Figure 1B). Isolates from Italy (MT012098) and Brazil (MT126808) clustered with different USA strains (Figure 1A & B). Notably, isolates from same country or geographical location formed a mosaic pattern of phylogenetic placements of countries’ isolates. For viral transmission, contact between the individuals is also an important factor, supposedly due to which the spread of identical strains across the border of neighboring countries is more likely. But we obtained a pattern where Indian strains showed highest similarity with USA and China strains, Australian strains with USA strains, Italy and Brazilian strains with strains isolated from USA among others. This depicts the viral spread across different communities. However, as genomes of SARS-CoV-2 were available mostly from USA and China, sampling biases is evident in analyzed dataset as available on NCBI. Thus, it is plausible for strains from other countries to show most similarity with strains from these two countries. In the near future as more and more genome sequences will become available from different geographical locations; more accurate patterns of their relatedness across the globe will become available

**Figure 1:**
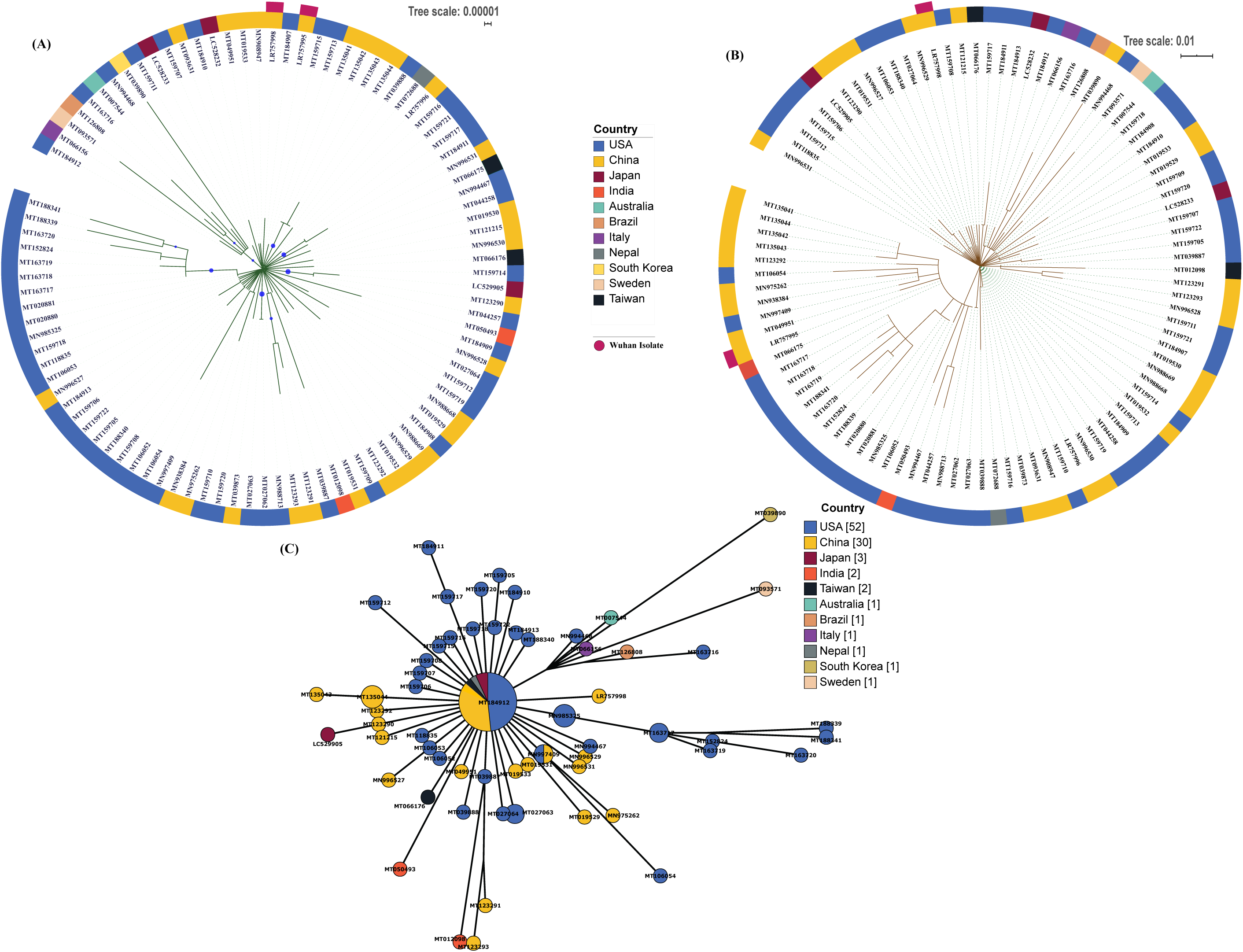
A) Core genome based phylogenetic analysis of SARS-CoV-2 isolates using the Maximum Likelihood method based on the Tamura-Nei model. The analysis involved 95 SARS-CoV-2 sequences with a total of 28451 nucleotide positions. Bootstrap values more than 70% are shown on branches as blue dots with sizes corresponding to the bootstrap values. The coloured circle represents the country of origin of each isolate. The two isolates from Wuhan are marked separately on the outside of the ring. B) SNP based phylogeny of SARS-CoV-2 isolates. Highly similar genomes of coronaviruses were taken as input by Parsnp. Whole-genome alignments were made using libMUSCLE aligner using the annotated genome of MT121215 strain as reference. Parsnp identifies the maximal unique matches (MUMs) among the query genomes provided in a single directory. As only genomes within a specified MUMI distance threshold are recruited, option -c to force include all the strains was used. The output phylogeny based on Single nucleotide polymorphisms was obtained following variant calling on core-genome alignment. C) The minimum spanning tree generated using Maximum Likelihood method and Tamura-Nei model showing the genetic relationships of SARS-CoV-2 isolates with their geographical distribution.

### SNPs in the SARS-CoV-2 genomes

SNPs in all predicted ORFs in each genome were analyzed using SARS-CoV-2/SH01/human/2020/CHN as a reference. SNPs were determined using maximum unique matches between the genomes of coronavirus, we observed that the strains isolated from USA (MT188341; MN985325; MT020881; MT020880; MT163719; MT163718; MT163717; MT152824; MT163720; MT188339) are the most evolved and they carry set of unique point mutations (Table2) in nsp13, nsp14, nsp15, nsp16 (present in orf1b polyprotein region) and S-Protein. All the mutated proteins are non-structural proteins (NSP) functionally involved in forming viral replication-transcription complexes (RTC) [46]. For instance, non-structural protein 13 (nsp13), belongs to helicase superfamily 1 and is putatively involved in viral RNA replication through RNA-DNA duplex unwinding [47] whereas nsp14 and nsp15 are exoribonuclease and endoribonuclease, respectively [48, 49]. nsp16 functions as a mRNA cap-1 methyltransferase [50]. All these proteins containing SNPs at several positions (Table 2) indicate that viral machinery for its RNA replication and processing is utmost evolved in strains from USA as compared to the other countries. Further, we analyzed the SNPs at protein level and interestingly in ORF1b protein, there were amino acid substitutions at P1327L, Y1364C and S2540F in USA isolates. One isolate namely USA0/MN1-MDH1/2020 (MT188341) carried amino-acid addition at 2540 position leading to shift in amino acid frame their onwards (Figure 2), which might affect the functioning of nsp16 (2′-O-MTase). But no changes were observed in Indian isolates, thus found similar to Chinese isolate. As the proteins involved in viral replication are evolving rapidly, this highlights the need to consider these mutants in order to develop the treatment strategies.

**Table 2:**
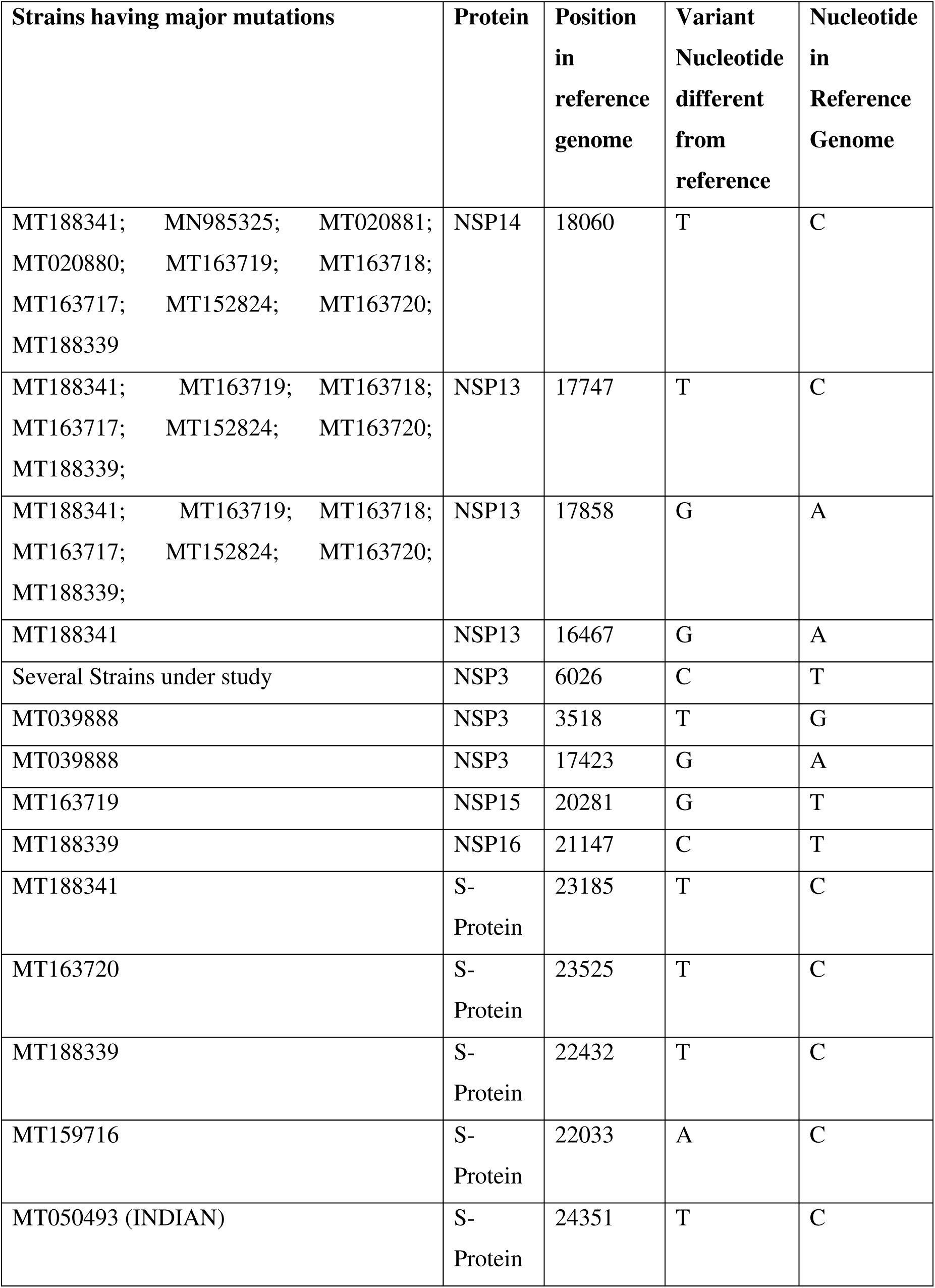
Major mutations present in different isolates of SARS-CoV-2 at different locations.

| **Strains having major mutations** | **Protein** | **Position**<br>**in**<br>**reference**<br>**genome** | **Variant**<br>**Nucleotide**<br>**different**<br>**from**<br>**reference** | **Nucleotide**<br>**in**<br>**Reference**<br>**Genome** |
| --- | --- | --- | --- | --- |
| MT188341; MN985325; MT020881;<br>MT020880; MT163719; MT163718;<br>MT163717; MT152824; MT163720;<br>MT188339 | NSP14 | 18060 | T | C |
| MT188341; MT163719; MT163718;<br>MT163717; MT152824; MT163720;<br>MT188339; | NSP13 | 17747 | T | C |
| MT188341; MT163719; MT163718;<br>MT163717; MT152824; MT163720;<br>MT188339; | NSP13 | 17858 | G | A |
| MT188341 | NSP13 | 16467 | G | A |
| Several Strains under study | NSP3 | 6026 | C | T |
| MT039888 | NSP3 | 3518 | T | G |
| MT039888 | NSP3 | 17423 | G | A |
| MT163719 | NSP15 | 20281 | G | T |
| MT188339 | NSP16 | 21147 | C | T |
| MT188341 | S-Protein | 23185 | T | C |
| MT163720 | S-Protein | 23525 | T | C |
| MT188339 | S-Protein | 22432 | T | C |
| MT159716 | S-Protein | 22033 | A | C |
| MT050493 (INDIAN) | S-Protein | 24351 | T | C |

**Figure 2:**
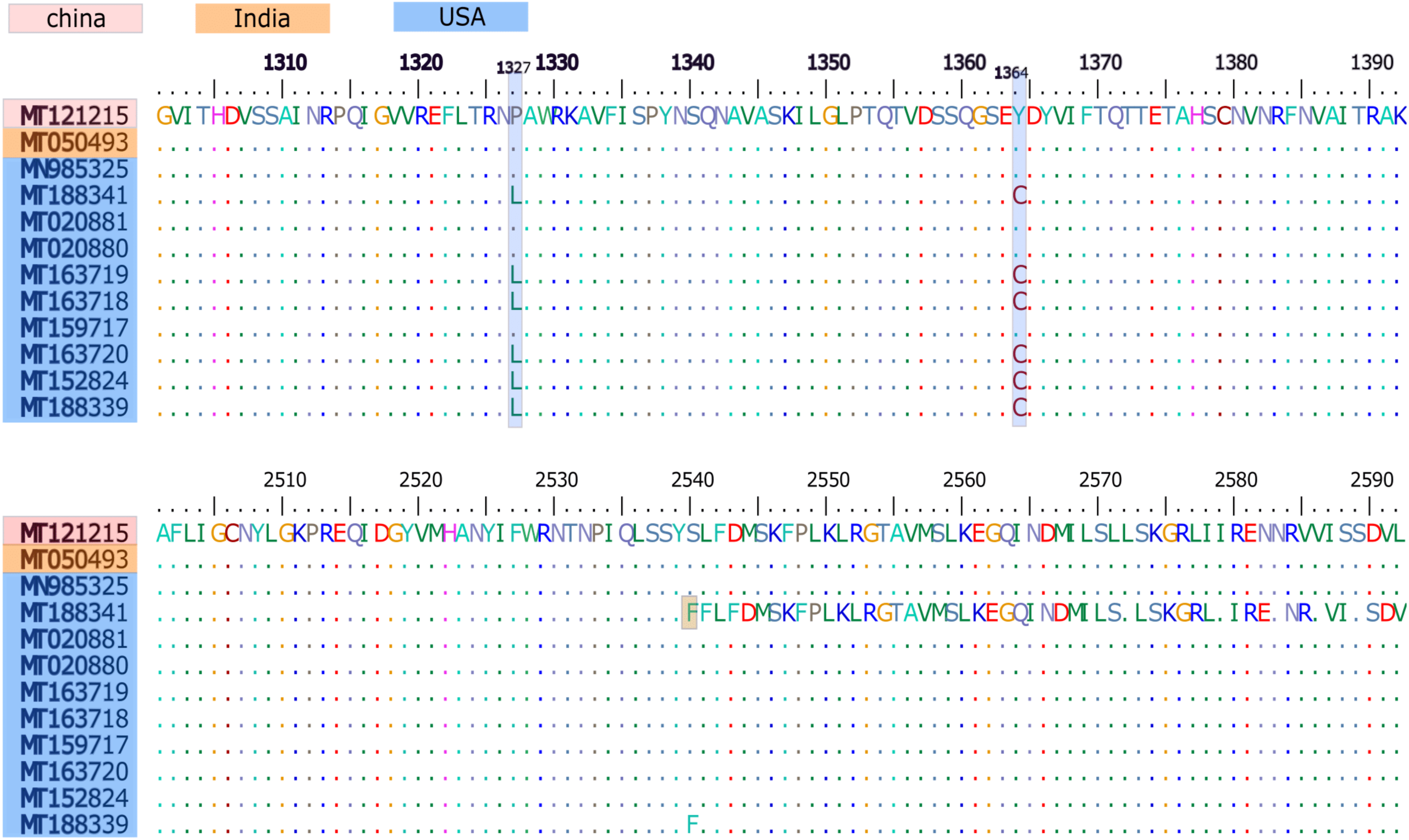
Multiple sequence alignment of ORF1b protein showing amino acid substitutions at three positions: P1327L, Y1364C and S2540F. The isolate USA/MN1-MDH1/2020 (MT188341) showed an amino-acid addition leading to change in amino acid frame from position 2540 onwards.

### Direction of selection of SARS-CoV-2 genes

Our analysis revealed that ORF8 (121 a.a.) (dN/dS= 35.8) along with ORF3a (275 bp) (dN/dS= 8.95) showed highest dN/dS values among the nine ORFs thus, have much greater number of non-synonymous substitutions than the synonymous substitution (Figure 3D). Values of dN/dS >>1 are indicative of strong divergent lineage [51]. Thus, both of these proteins are evolving under high selection pressure and are highly divergent ORFs across strains. Two other proteins, ORF1ab polyprotein (dN/dS= 0.996, 0.575) and S protein (dN/dS= 0.88) might confer selective advantage with host challenges and survival. The dN/dS rates nearly 1 and greater than 1 suggests that the strains are coping up with the challenges *i.e.*, immune responses and inhibitory environment of host cells [52]. The other gene clusters namely M-protein and orf1a polyprotein did not possess at least three unique sequences necessary for the analysis, hence, they should be similar across the strains. The two genes ORF1ab polyprotein encodes for protein translation and post translation modification found to be evolved which actively translates, enhance the multiplication and facilitates growth of virus inside the host. Similarly, the S protein which helps in the entry of virus to the host cells by surpassing the cell membrane found to be accelerated towards positive selection confirming the successful ability of enzyme to initiate the infection. Another positive diversifying gene N protein encodes for nucleocapsid formation which protects the genetic material of virus form host immune responses such as cellular proteases. Overall, the data represent that the growth and multiplication related genes are highly evolving. The other proteins with dN/dS values equal to zero suggesting a conserved repertoire of genes.

**Figure 3:**
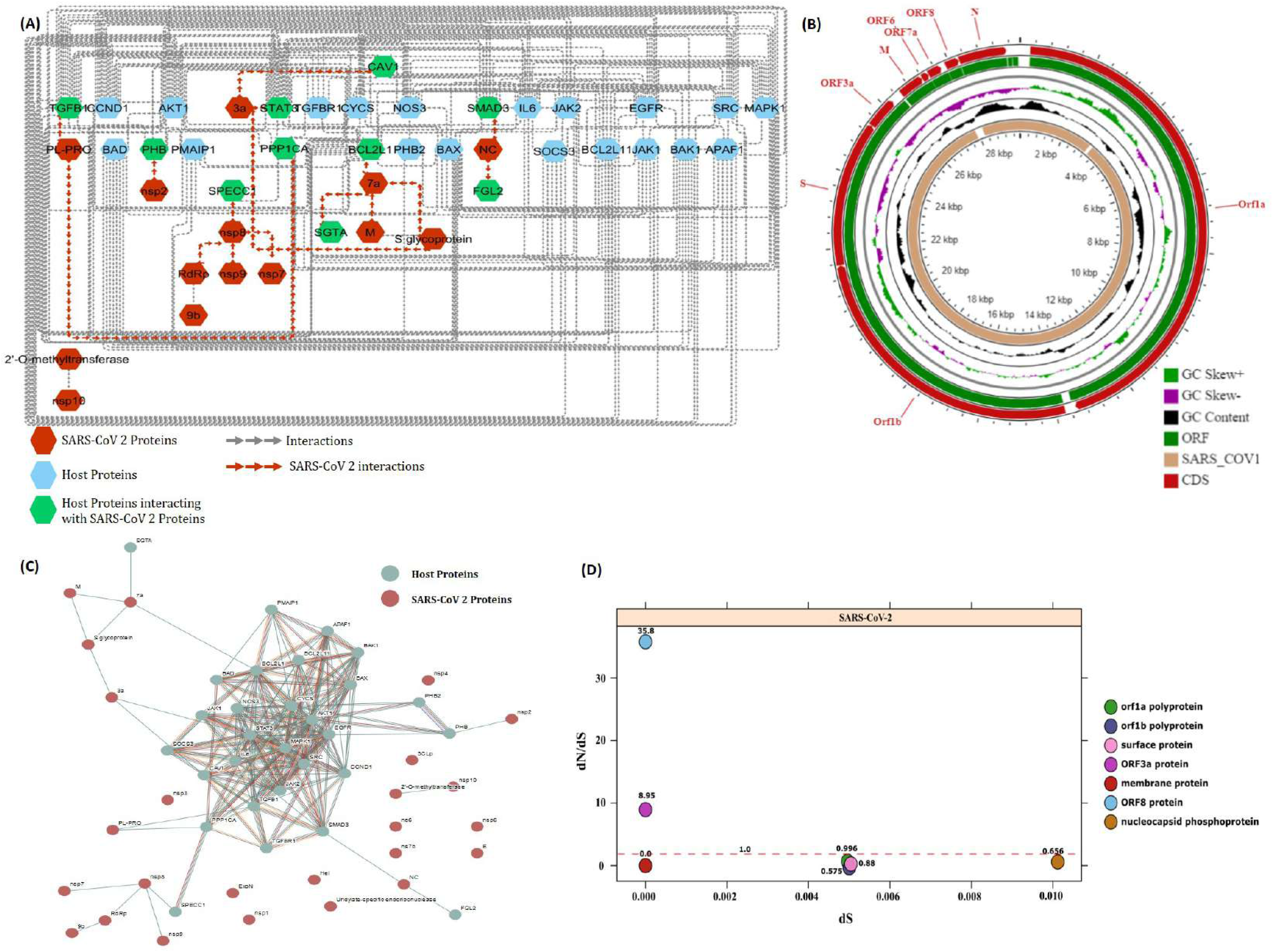
(A) SARS-CoV-2 -Host interactome analysis. Sub-set network highlighting SARS-CoV-2 and host nodes targeting each other. In total, nine direct interactions were observed (shown with red arrows). (B) Circular genome map of SARS-CoV-2 with genome size of 29.8 Kb generated using CGView. The genome of SARS-CoV 2 is also compared with that of SARS-CoV genome. The ruler for genome size is shown as innermost ring where Kbp stands for kilo base pairs. Concentric circles from inside to outside denote: SARS-CoV genome (used as reference), G + C content, G + C skew, predicted ORFs in SARS-CoV-2 genome and annotated CDS in SARS-CoV-2 genome. Gaps in alignment are shown in white. The positive and negative deviation from mean G + C content and G + C skew are represented with outward and inward peaks respectively. (C) SARS-CoV 2 and Host interactome generated using Virus.STRING interaction database v10.5. Both interacting and non-interacting viral proteins are shown. (D) Estimation of purifying natural selection pressure in nine coding sequences of SARS-CoV-2. dN/dS values are plotted as a function of dS.

### SARS-CoV-2-Host interactome unveils immunopathogenesis of COVID-19

Although the primary mode of infection is human to human transmission through close contact, which occurs via spraying of nasal droplets from the infected person, yet the primary site of infection and pathogenesis of SARS-CoV-2 is still not clear and under investigation. To explore the role of SARS-CoV-2 proteins in host immune evasion, the SARSCoV-2 proteins were mapped over host proteome database (Figure 3B & Table 3). We identified a total of 28 proteins from host proteome forming close association with 25 viral proteins present in 9 ORFs of SARS-CoV-2 (Figure 3C). The network was trimmed in Cytoscape v3.7.2 where only interacting proteins were selected. Only 12 viral proteins were found to interact with host proteins (Figure 3A). Detailed analysis of interactome highlighted 9 host proteins in direct association with 6 viral proteins. Further, the network was analyzed for identification of regulatory hubs based on degree analysis. We identified mitogen activated protein kinase 1 (MAPK1) and AKT proteins as major hubs forming 24 and 21 interactions in the network respectively, highlighting their crucial role in pathogenesis. Recently, Huang *et al*, demonstrated the role of Mitogen activated protein kinase (MAPK) in COVID-19 mediated blood immune responses in infected patients [53] and showed that MAPK activation certainly plays a major defense mechanism.

**Table 3:**
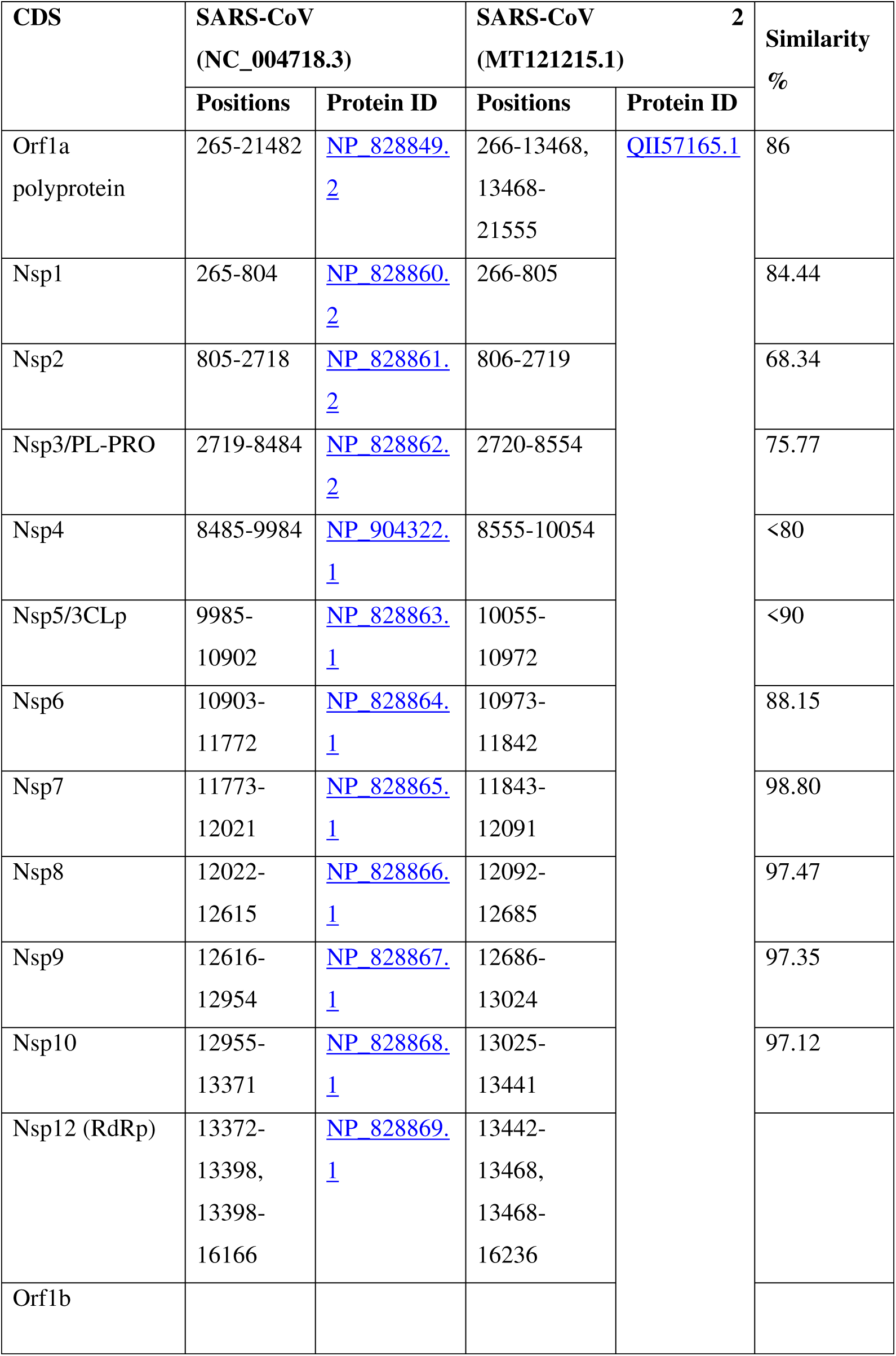
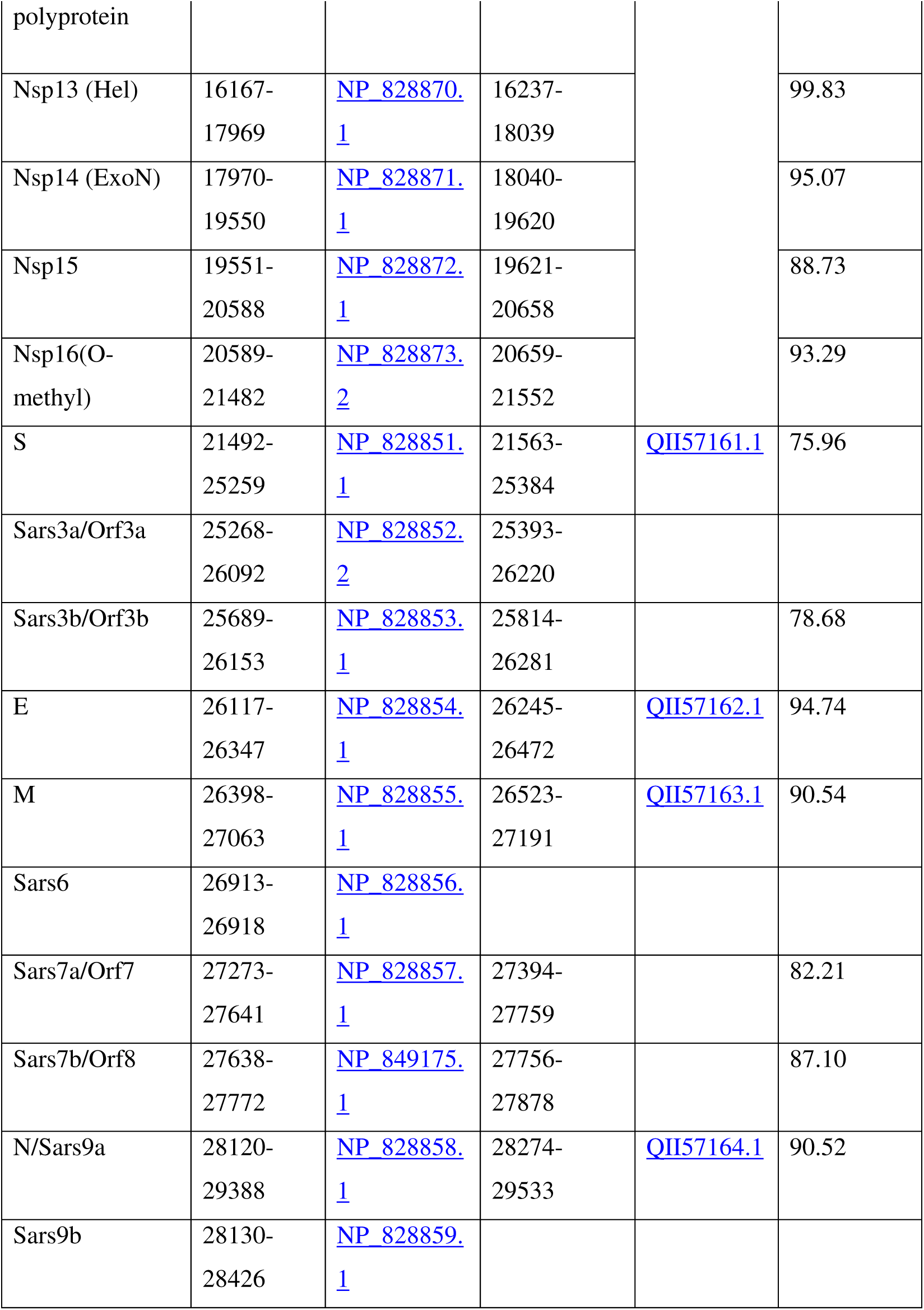
Description of SARS-CoV2 proteins and its similarity in comparison to SARS-CoV used for PPI prediction.

Gene Ontology based functional annotation studies predicted the role of direct interactions of several viral proteins with host proteins. One such protein is non-structural protein2 (nsp2) which directly interacts with host Prohibitin (PHB), a known regulator of cell proliferation and maintains functional integrity of mitochondria [54]. SARS-CoV nsp2 is also known for its interaction with host PHB1 and PHB2 [55]. Nsp2 is a methyltransferase like domain that is known to mediate mRNA cap 2’-O-ribose methylation to the 5’-cap structure of viral mRNAs. This N7-methylguanosine cap is required for the action of nsp16 (2’-O-methyltransferase) and nsp10 complex [56]. This 5’-capping of viral RNA plays a crucial role in escape of virus from innate immunity recognition [56]. Hence, nsp2 -is responsible for modulating host cell survival strategies by altering host cell environment [55]. Based on network predicted we propose nsp16/nsp10 interface as a better drug target for anti-coronavirus drugs corresponding to the prediction made by Chen and group (2011) [56].

Similarly, the viral protein Papain-like proteinase (PL-PRO) which has deubiquitinase and deISGylating activity is responsible for cleaving viral polyprotein into 3 mature proteins which are essential for viral replication [57]. Our study showed that PL-PRO directly interacts with PPP1CA which is a protein phosphatase that associates with over 200 regulatory host proteins to form highly specific holoenzymes. PL-PRO is also found to interact with TGFβ which is a beta transforming growth factor and promotes T-helper 17 cells (Th17) and regulatory T-cells (T*reg*) differentiation [58]. Reports have shown the PL-PRO induced upregulation of TGFβ in human promonocytes via MAPK pathway result in pro-fibrotic responses [59]. This reflects that viral PL-PRO antagonises innate immune system and is directly involved in the pathogenicity of SARS-CoV-2 induced pulmonary fibrosis [56, 58]. Many COVID-19 patients develop acute respiratory distress syndrome (ADRS) which leads to pulmonary edema and lung failure [60, 61]. These symptoms are because of cytokine storm manifesting elevated levels of pro-inflammatory cytokines like IL6, IFNγ, IL17, IL1β etc [61]. These results are in agreement with our prediction where we found IL6 as an interacting partner. Our study also showed JAK1/2 as an interacting partner which is known for IFNγ signaling. It is well known that TGFβ along with IL6 and STAT3 promotes Th17 differentiation by inhibiting SOCS3 [62]. Th17 is a source of IL17, which is commonly found in serum samples of COVID-19 patients [61, 63]. Hence, our interactome is supported from these findings where we found SOCS3, STAT3, JAK1/2 as an interacting partner [64]. The results suggested that proinflammatory cytokine storm is one of the reasons for SARS-CoV-2 mediated immunopathogenesis.

In the next cycle of physical events the viral protein NC (nucleoprotein), which is a major structural part of SARV family associates with the genomic RNA to form a flexible, helical nucleocapsid. Interaction of this protein with SMAD3 leads to inhibition of apoptosis of SARS-CoV infected lung cells [65], which is a successful strategy of immune evasion by the virus. More complex and multiple associations of ORF7a viral protein which is a non-structural protein and known as growth factor for SARS family viruses, directly captures BCL2L1 which is a potent regulator of apoptosis. Tan *et al.* (2007) have shown that SARS-CoV ORF7a protein induces apoptosis by interacting with Bcl XL protein which is responsible for lymphopenia, an abnormality found in SARS-CoV infected patients [66]. Another target of viral ORF7a protein is SGTA (Small glutamine-rich tetratricopeptide repeat) which is an ATPase regulator and promotes viral encapsulation [67]. Subordinate viral proteins M (Membrane), S (Glycoprotein) and ORF3a (viroporin) were found to interact with each other. This interaction is important for viral cell formation and budding [68, 69]. Studies have shown the localization of ORF3a protein in Golgi apparatus of SARS-CoV infected patients along with M protein and responsible for viral budding and cell injury [70]. ORF3a protein also targets the functioning of CAV1 (Caveolin 1), caveolae protein, acts as a scaffolding protein within caveolar membranes. CAV1 has been reported to be involved in viral replication, persistence, and the potential role in pathogenesis in HIV infection also [71]. Thus, ORF3a interactions will upregulate viral replication thus playing a very crucial role in pathogenesis. Multiple methyltransferase assembly viral proteins (nsp7, nsp8, nsp9, RdRp) which are nuclear structural proteins were observed to target the SPECC1 proteins and linked with cytokinesis and spindle formations during division. Thus, major viral assembly also targets the proteins linked with immunity and cell division. Taken together, we estimated that SARS-CoV-2 manipulate multiple host proteins for its survival while, their interaction is also a reason for immunopathogenesis.

### Conclusions

As COVID-19 continues to impact virtually all human lives worldwide due to its extremely contagious nature, it has spiked the interest of scientific community all over the world to understand better the pathogenesis of the novel SARS-CoV-2. In this study, the analysis was performed on the genomes of the novel SARS-CoV-2 isolates recently reported from different countries to understand viral pathogenesis. With the limited data available so far, we observed no direct transmission pattern of the novel SARS-CoV-2 in the neighboring countries through our analyses of the phylogenomic relatedness of geographical isolates. The isolates from same locations were phylogenetically distant, for instance, isolates from the USA and China. Thus, there appears to be a mosaic pattern of transmission indicative of the result of infected human travel across different countries. As COVID-19 transited from epidemic to pandemic within a short time, it does not look surprising from the genome structures of the viral isolates. The genomes of six isolates, specifically from the USA, were found to harbor unique amino acid SNPs and showed amino acid substitutions in ORF1b protein and S-protein, while one of them also harbored an amino-acid addition. This is suggestive of the severity of the mutating viral genomes within the population of the USA. These proteins are directly involved in the formation of viral replication-transcription complexes (RTC). Therefore, we argue that the novel SARS-CoV-2 has fast evolving replicative machinery and that it is urgent to consider these mutants to develop strategies for COVID-19 treatment. The ORF1ab polyprotein protein and S-protein were also found to have dN/dS values approaching 1 and thus might confer a selective advantage to evade host responsive mechanisms. The construction of SARS-CoV-2-human interactome revealed that its pathogenicity is mediated by a surge in pro-inflammatory cytokine. It is predicted that major immune-pathogenicity mechanism by SARS-CoV-2 includes the host cell environment alteration by disintegration by signal transduction pathways and immunity evasion by several protection mechanisms. The mode of entry of this virus by S-proteins inside the host cell is still unclear but it might be similar to SARS CoV-1 like viruses. Lastly, we believe as more data accumulate for COVID-19 the evolutionary pattern will become much clear.

## Authors Contribution

RL, RK, HV, VG, US conceived and designed the study. RK, HV, NS, US, VG, MS, SN, PH executed the analysis and prepared figures. RK, HV, RK, NS, US, VG, MS, SN, PH, CT, NN, SA, CDR, MV wrote the manuscript with contributions from all authors. YS and RKN provided time to time guidance.

## Acknowledgements

RK acknowledges Magadh University, Bodh Gaya for providing support. RL and US also acknowledge The National Academy of Sciences, India, for support under the NASI-Senior Scientist Platinum Jubilee Fellowship Scheme. NS, SN, CT acknowledge Council of Scientific and Industrial Research (CSIR), New Delhi for doctoral fellowships. HV would like to thank Ramjas College, University of Delhi, Delhi for providing support. VG and MS acknowledge Phixgen Pvt. Ltd. for research fellowship. PH would like to thank Maitreyi College, University of Delhi, Delhi for providing support.

